# Could a Neuroscientist Understand a Microprocessor?

**DOI:** 10.1101/055624

**Authors:** Eric Jonas, Konrad Paul Kording

## Abstract

There is a popular belief in neuroscience that we are primarily data limited, and that producing large, multimodal, and complex datasets will, with the help of advanced data analysis algorithms, lead to fundamental insights into the way the brain processes information. These datasets do not yet exist, and if they did we would have no way of evaluating whether or not the algorithmically-generated insights were sufficient or even correct. To address this, here we take a classical microprocessor as a model organism, and use our ability to perform arbitrary experiments on it to see if popular data analysis methods from neuroscience can elucidate the way it processes information. Microprocessors are among those artificial information processing systems that are both complex and that we understand at all levels, from the overall logical flow, via logical gates, to the dynamics of transistors. We show that the approaches reveal interesting structure in the data but do not meaningfully describe the hierarchy of information processing in the microprocessor. This suggests current analytic approaches in neuroscience may fall short of producing meaningful understanding of neural systems, regardless of the amount of data. Additionally, we argue for scientists using complex non-linear dynamical systems with known ground truth, such as the microprocessor as a validation platform for time-series and structure discovery methods.

**Author Summary:** Neuroscience is held back by the fact that it is hard to evaluate if a conclusion is correct; the complexity of the systems under study and their experimental inaccessability make the assessment of algorithmic and data analytic technqiues challenging at best. We thus argue for testing approaches using known artifacts, where the correct interpretation is known. Here we present a microprocessor platform as one such test case. We find that many approaches in neuroscience, when used na•vely, fall short of producing a meaningful understanding.

## Introduction

The development of high-throughput techniques for studying neural systems is bringing about an era of big-data neuroscience [1, 2]. Scientists are beginning to reconstruct connectivity [3], record activity [4], and simulate computation [5] at unprecedented scales. However, even state-of-the-art neuroscientific studies are still quite limited in organism complexity and spatiotemporal resolution [?, 6, 7]. It is hard to evaluate how much scaling these techniques will help us understand the brain.

In neuroscience it can be difficult to evaluate the quality of a particular model or analysis method, especially in the absence of known truth. However, there are other systems, in particular man made ones that we do understand. As such, one can take a human-engineered system and ask if the methods used for studying biological systems would allow understanding the artificial system. In this way, we take as inspiration Yuri Lazbnick's well-known 2002 critique of modeling in molecular biology, “Could a biologist fix a radio?” [8]. However, a radio is clearly much simpler than the nervous system, leading us to seek out a more complex, yet still well-understood engineered system. The microprocessors in early computing systems can serve this function.

Here we will try to understand a known artificial system, a classical microprocessor by applying data analysis methods from neuroscience. We want to see what kind of an understanding would emerge from using a broad range of currently popular data analysis methods. To do so, we will analyze the connections on the chip, the effects of destroying individual transistors, single-unit tuning curves, the joint statistics across transistors, local activities, estimated connections, and whole-device recordings. For each of these, we will use standard techniques that are popular in the field of neuroscience. We find that many measures are surprisingly similar between the brain and the processor but that our results do not lead to a meaningful understanding of the processor. The analysis can not produce the hierarchical understanding of information processing that most students of electrical engineering obtain. It suggests that the availability of unlimited data, as we have for the processor, is in no way sufficient to allow a real understanding of the brain. We argue that when studying a complex system like the brain, methods and approaches should first be sanity checked on complex man-made systems that share many of the violations of modeling assumptions of the real system.

## An engineered model organism

The MOS 6502 (and the virtually identical MOS 6507) were the processors in the Apple I, the Commodore 64, and the Atari Video Game System (VCS) (see [9] for a comprehensive review). The Visual6502 team reverse-engineered the 6507 from physical integrated circuits [10] by chemically removing the epoxy layer and imaging the silicon die with a light microscope. Much like with current connectomics work [11, 12], a combination of algorithmic and human-based approaches were used to label regions, identify circuit structures, and ultimately produce a transistor-accurate netlist (a full connectome) for this processor consisting of 3510 enhancement-mode transistors. Several other support chips, including the Television Interface Adaptor (TIA) were also reverse-engineered and a cycle-accurate simulator was written that can simulate the voltage on every wire and the state of every transistor. The reconstruction has sufficient fidelity to run a variety of classic video games, which we will detail below. The simulation generates roughly 1.5GB/sec of state information, allowing a real big-data analysis of the processor.

The simplicity of early video games has led to their use as model systems for reinforcement learning [13] and computational complexity research [14]. The video game system (“whole animal”) has a well defined output in each of the three behavioral conditions (games). It produces an input-dependent output that is dynamic, and, in the opinion of the authors, quite exciting. It can be seen as a more complex version of the Mus Silicium project [15]. It is also a concrete implementation of a thought experiment that has been mentioned on and off in the literature [16–19]. The richness of the dynamics and our knowledge about its inner workings makes it an attractive test case for approaches in neuroscience.

**Fig. 1.**
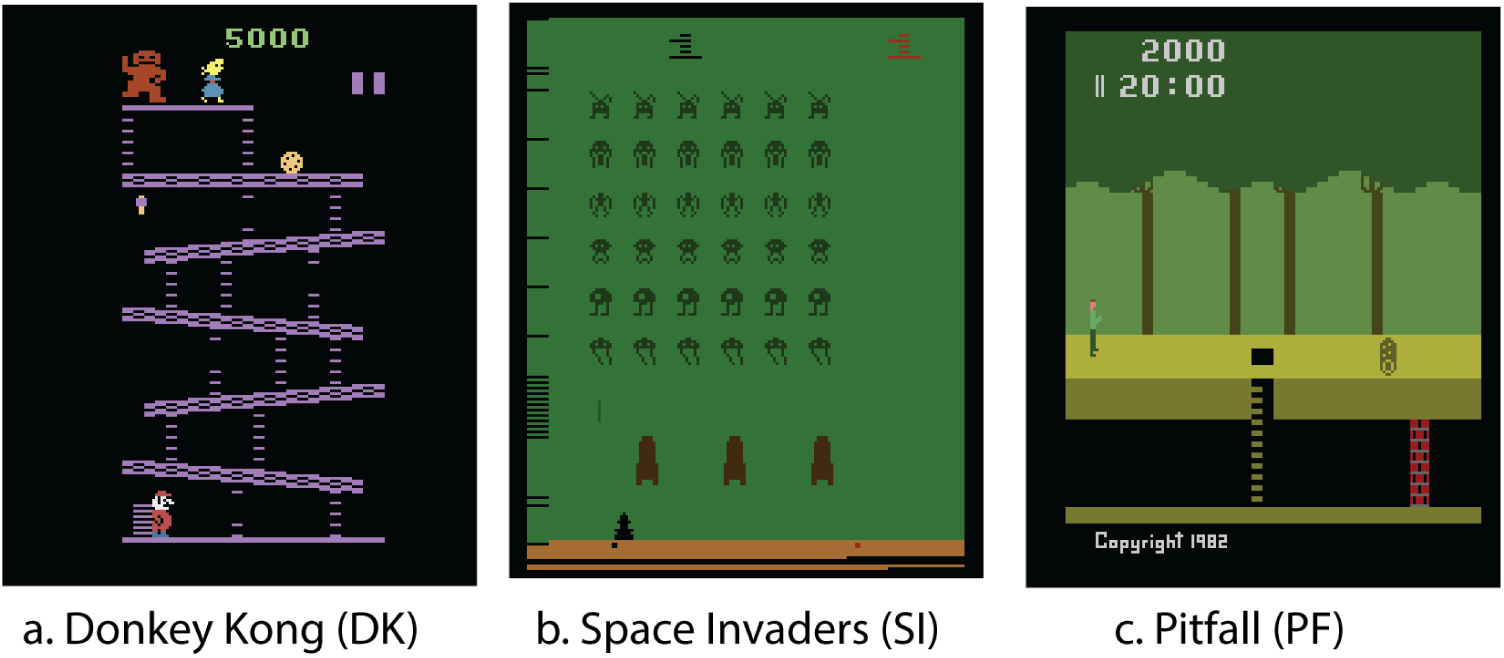
Example behaviors. We use three classical video games as example behaviors for our model organism – (**A**) Donkey Kong (1981), (**B**) Space Invaders (1978), and (**C**) Pitfall (1981).

For this paper we will only use three behaviors, three different games. Obviously these “behaviors” are qualitatively different from those of animals and may seem more complicated. However, even the simple behaviors that are studied in neuroscience still involve a plethora of components, typically including the allocation of attention, cognitive processing, and multiple modalities of inputs and outputs. As such, the breadth of ongoing computation in the processor may actually be simpler than those in the brain.

The objective of clever experimental design in neuroscience often is to find behaviors that only engage one kind of computation in the brain. In the same way, all our experiments on the chip will be limited by us only using these games to probe it. As much as more neuroscience is interested in naturalistic behaviors [20], here we analyze a naturalistic behavior of the chip. In the future it may be possible to excute simpler, custom code on the processor to tease apart aspects of computation, but we currently lack such capability in biological organisms.

Much has been written about the differences between computation in silico and computation in vivo [21, 22]—the stochasticity, redundancy, and robustness [23] present in biological systems seems dramatically different from that of a microprocessor. But there are many parallels we can draw between the two types of systems. Both systems consist of interconnections of a large number of simpler, stereotyped computing units. They operate on multiple timescales. They consist of somewhat specialized modules organized hierarchically. They can flexibly route information and retain memory over time. Despite many differences there are also many similarities. We do not wish to overstate this case – in many ways, the functional specialization present in a large mammalian brain far eclipses that present in the processor. Indeed, the processor's scale and specialization share more in common with *C. elegans* than a mouse.

Yet many of the differences should make analysing the chip easier than analyzing the brain. For example, it has a clearer architecture and far fewer modules. The human brain has hundreds of different types of neurons and a similar diversity of proteins at each individual synapse [24], whereas our model microprocessor has only one type of transistor (which has only three terminals). The processor is deterministic while neurons exhibit various sources of randomness. With just a couple thousand transistors it is also far smaller. And, above all, in the simulation it is fully accessible to any and all experimental manipulations that we might want to do on it.

**Fig 2.**
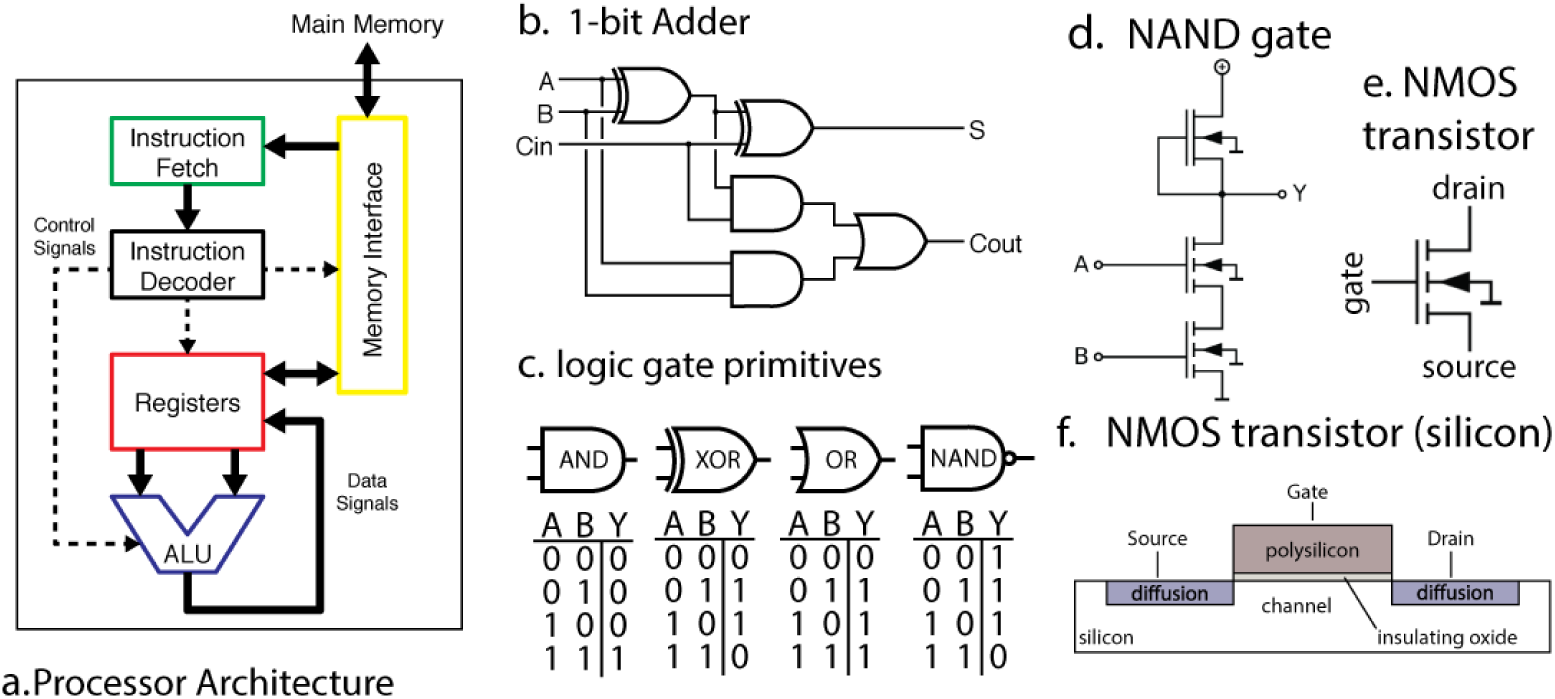
A microprocessor is understood at all levels. (**A**) The instruction fetcher obtains the next instruction from memory. This then gets converted into electrical signals by the instruction decoder, and these signals enable and disable various internal parts of the processor, such as registers and the arithmetic logic unit (ALU). The ALU performs mathematical operations such as addition and subtraction. The results of these computations can then be written back to the registers or memory. (**B**) Within the ALU there are well-known circuits, such as this one-bit adder, which sums two one-bit signals and computes the result and a carry signal. (**C**) Each logic gate in (**B**) has a known truth table and is implemented by a small number of transistors. (**D**) A single NAND gate is comprised of transistors, each transistor having three terminals (**E**). We know (**F**) the precise silicon layout of each transistor.

## What does it mean to understand a system

Importantly, the processor allows us to ask “do we really understand this system?” Most scientists have at least behavioral-level experience with these classical video game systems, and many in our community, including some electrophysiologists and computational neuroscientists, have formal training in computer science, electrical engineering, computer architecture, and software engineering. As such, we believe that most neuroscientists may have better intuitions about the workings of a processor than about the workings of the brain.

What constitutes an understanding of a system? Lazbnick's original paper argued that understanding was achieved when one could “fix” a broken implementation. Understanding of a particular region or part of a system would occur when one could describe so accurately the inputs, the transformation, and the outputs that one brain region could be replaced with an entirely synthetic component. Indeed, some neuroengineers are following this path for sensory [25] and memory [26] systems. Alternatively, we could seek to understand a system at differing, complementary levels of analysis, as David Marr and Tomaso Poggio outlined in 1982 [27]. First, we can ask if we understand what the system does at the computational level: what is the problem it is seeking to solve via computation? We can ask how the system performs this task algorithmically : what processes does it employ to manipulate internal representations? Finally, we can seek to understand how the system implements the above algorithms at a physical level. What are the characteristics of the underlying implementation (in the case of neurons, ion channels, synaptic conductances, neural connectivity, and so on) that give rise to the execution of the algorithm? Ultimately, we want to understand the brain at all these levels.

In this paper, much as in systems neuroscience, we consider the quest to gain an understanding of how circuit elements give rise to computation. Computer architecture studies how small circuit elements, like registers and adders, give rise to a system capable of performing general-purpose computation. When it comes to the processor, we understand this level extremely well, as it is taught to most computer science undergraduates. Knowing what a satisfying answer to “how does a processor compute?” looks like makes it easy to evaluate how much we learn from an experiment or an analysis.

## What would a satisfying understanding of the processor look like?

We can draw from our understanding of computer architecture to firmly ground what a full understanding of a processor would look like (fig 2). The processor is used to implement a computing machine. It implements a finite state machine which sequentially reads in an instruction from memory (fig 2a, green) and then either modifies its internal state or interacts with the world. The internal state is stored in a collection of byte-wide registers (fig 2a, red). As an example, the processor might read an instruction from memory telling it to add the contents of register A to the contents of register B. It then decodes this instruction, enabling the arithmetic logic unit (ALU, fig 2a, blue) to add those registers, storing the output. Optionally, the next instruction might save the result back out to RAM (fig 2a, yellow). It is this repeated cycle that gives rise to the complex series of behaviors we can observe in this system. Note that this description in many ways ignores the functions of the individual transistors, focusing instead on circuits modules like “registers” which are composed of many transistors, much as a systems neuroscientist might focus on a cytoarchitecturally-distinct area like hipppocampus as opposed to individual neurons.

Each of the functions within the processor contains algorithms and a specific implementation. Within the arithmetic logic unit, there is a byte wide adder, which is in part made of binary adders (fig 2b), which are made out of AND/NAND gates, which are made of transistors. This is in a similar way as the brain consists of regions, circuits, microcircuits, neurons, and synapses.

If we were to analyze a processor using techniques from systems neuroscience we would hope that it helps guide us towards the descriptions that we used above. In the rest of the paper we will apply neuroscience techniques to data from the processor. We will finally discuss how neuroscience can work towards techniques that will make real progress at moving us closer to a satisfying understanding of computation, in the chip, and in our brains.

## Results

Validating our understanding of complex systems is incredibly difficult when we do not know the actual ground truth. Thus we use an engineered system, the MOS6502, where we understand every aspect of its behavior at many levels. We will examine the processor at increasingly-fine spatial and temporal resolutions, eventually achieving true “big-data” scale : a “processor activity map”, with every transistor state and every wire voltage. As we apply the various techniques that are currently used in neuroscience we will ask how the analyses bring us closer to an understanding of the microprocessor (Fig. 2). We will use this well defined comparison to ask questions about the validity of current approaches to studying information processing in the brain.

**Fig 3.**
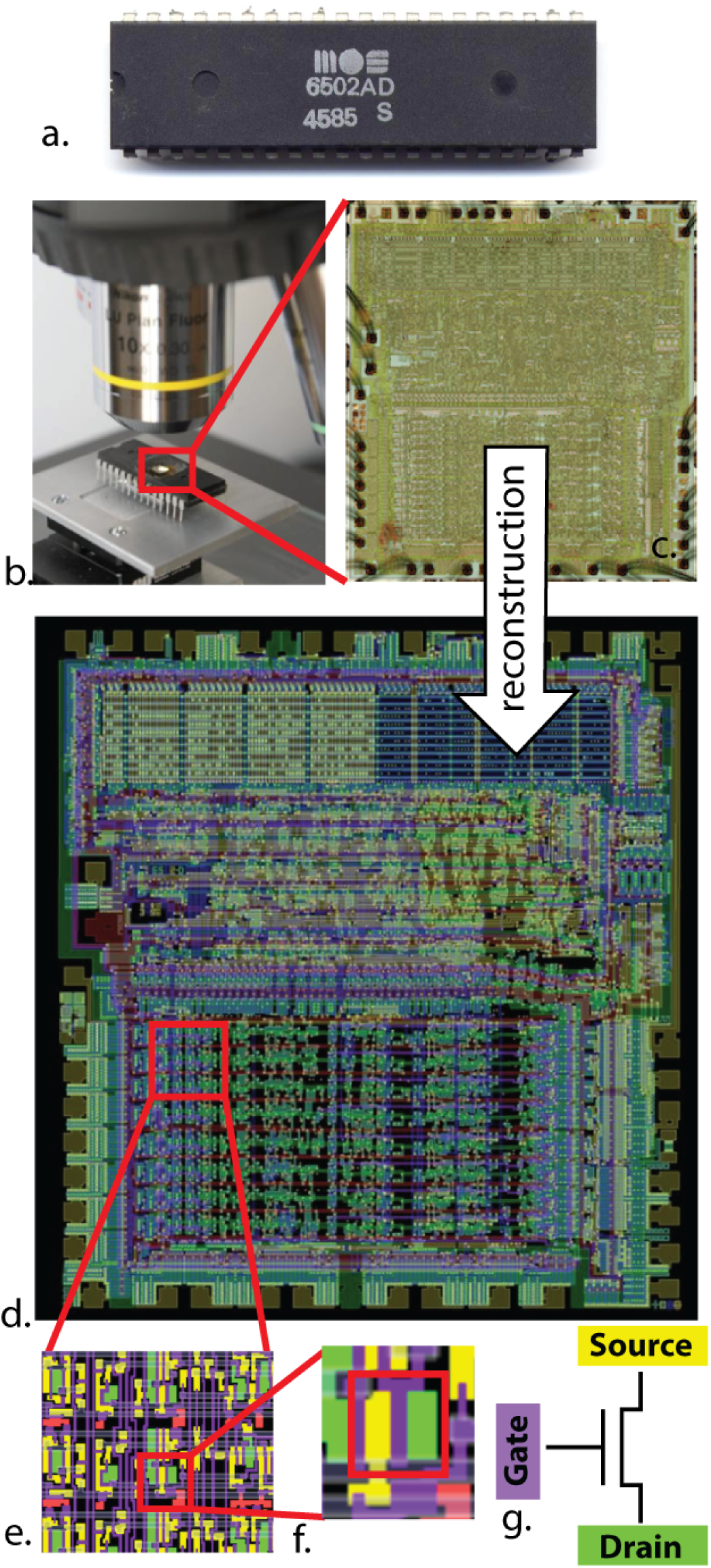
Optical reconstruction of the microprocessor to obtain its connectome. In [10], the (**A**) MOS 6502 silicon die was examined under a visible light microscope (**B**) to build up an image mosaic (**C**) of the chip surface. Computer vision algorithms were used to identify metal and silicon regions (**E**) to detect transistors (**F**), (**G**) ultimately producing a complete accurate netlist of the processor (**D**)

**Fig 4.**
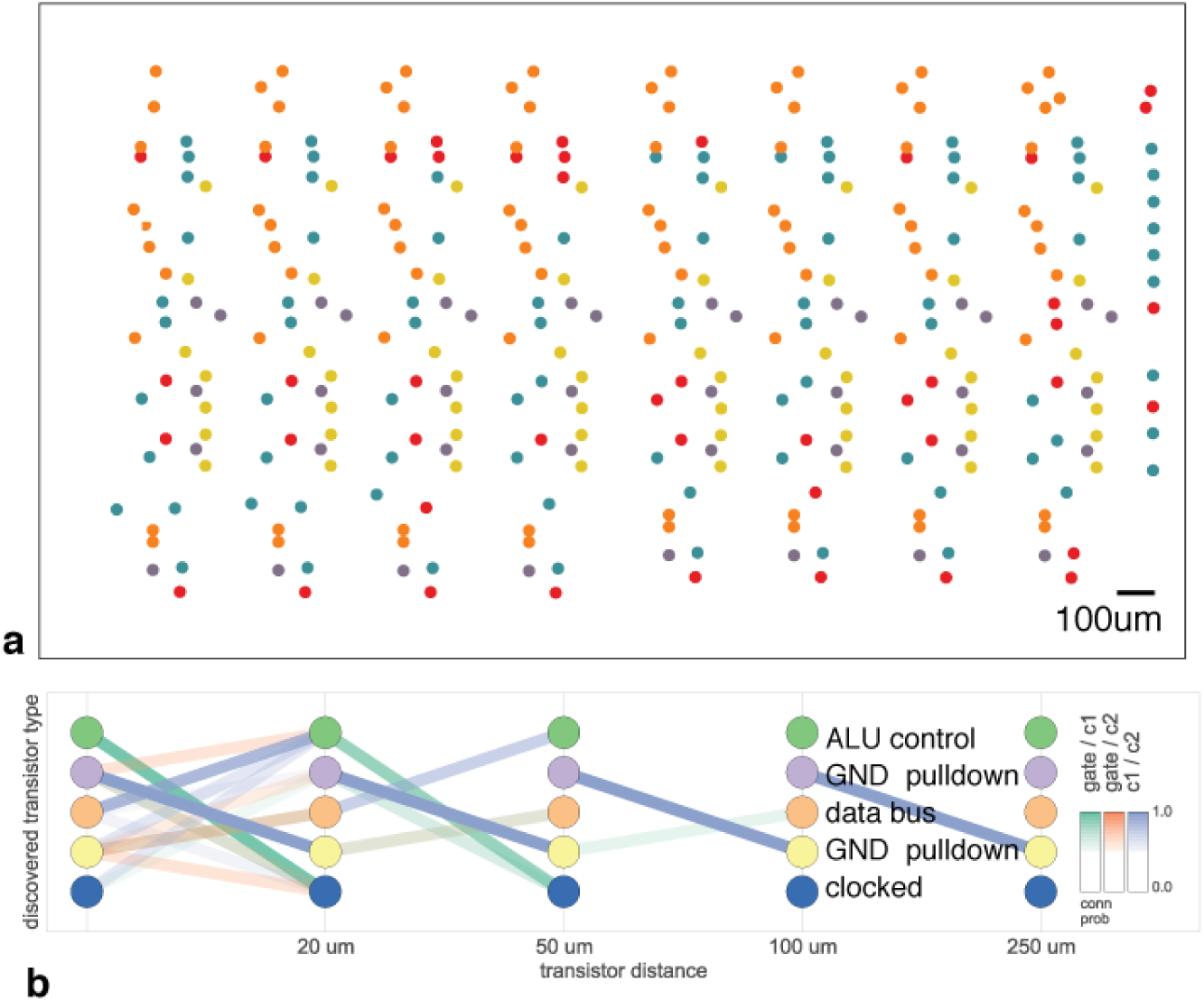
Discovering connectivity and cell type. Reproduced from [29]. (**A**) The spatial distribution of the transistors in each cluster show a clear pattern (**B**) The clusters and connectivity versus distance for connections between Gate and C1, Gate and C2, and C1 and C2 terminals on a transistor. Purple and yellow types have a terminal pulled down to ground and mostly function as inverters. The blue types are clocked, stateful transistors, green control the ALU and orange control the special data bus (SDB).

## Connectomics

The earliest investigations of neural systems were in-depth anatomical inquiries [28]. Fortunately, through large scale microscopy (Figure 3a) we have available the full 3d connectome of the system. In other words, we know how each transistor is connected to all the others. The reconstruction is so good, that we can now simulate this processor perfectly – indeed, were it not for the presence of the processor's connectome, this paper would not have been possible. This process is aided by the fact that we know a transistor's deterministic input-output function, whereas neurons are both stochastic and vastly more complex.

Recently several graph analysis methods ranging from classic [30] to modern [29, 31] approaches have been applied to neural connectomes. The approach in [29] was also applied to a region of this processor, attempting to identify both circuit motifs as well as transistor “types” (analogous to cell types) in the transistor wiring diagram. Figure 4 (adapted from [29]) shows the results of the analysis. We see that one identified transistor type contains the “clocked” transistors, which retain digital state. Two other types contain transistors with pins C1 or C2 connected to ground, mostly serving as inverters. An additional identified type controls the behavior of the three registers of interest (X, Y, and S) with respect to the SB data bus, either allowing them to latch or drive data from the bus. The repeat patterns of spatial connectivity are visible in Figure 4a, showing the man-made horizontal and vertical layout of the same types of transistors.

While superficially impressive, based on the results of these algorithms we still can not get anywhere near an understanding of the way the processor really works. Indeed, we know that for this processor there is only one physical “type” of transistor, and that the structure we recover is a complex combination of local and global circuitry.

In neuroscience, reconstructing all neurons and their connections perfectly is the dream of a large community studying connectomics [32, 33]. Current connectomics approaches are limited in their accuracy and ability to definitively identify synapses [12], Unfortunately, we do not yet have the techniques to also reconstruct the i/o function – neurotransmitter type, ion channel type, I/V curve of each synapse, etc. – of each neuron. But even if we did, just as in the case of the processor, we would face the problem of understanding the brain based on its connectome. As we do not have algorithms that go from anatomy to function at the moment that go considerably beyond cell-type clustering [29, 34, 35] it is far from obvious how a connectome would allow an understanding of the brain.

Note we are not suggesting connectomics is useless, quite the contrary – in the case of the processor the connectome was the first crucial step in enabling reliable, whole-brain-scale simulation. But even with the whole-brain connectome, extracting hierarchical organization and understanding the nature of the underlying computation is incredibly difficult.

## Lesion a single transistor at a time

Lesions studies allow us to study the causal effect of removing a part of the system. We thus chose a number of transistors and asked if they are necessary for each of the behaviors of the processor (figure 5. In other words, we asked if removed each transistor, if the processor would then still boot the game. Indeed, we found a subset of transistors that makes one of the behaviors (games) impossible. We can thus conclude they are uniquely necessary for the game – perhaps there is a Donkey Kong transistor or a Space Invaders transistor. Even if we can lesion each individual transistor, we do not get much closer to an understanding of how the processor really works.

This finding of course is grossly misleading. The transistors are not specific to any one behavior or game but rather implement simple functions, like full adders. The finding that some of them are important while others are not for a given game is only indirectly indicative of the transistor's role and is unlikely to generalize to other games. Lazebnik [8] made similar observations about this approach in molecular biology, suggesting biologists would obtain a large number of identical radios and shoot them with metal particles at short range, attempting to identify which damaged components gave rise to which broken phenotype.

This example nicely highlights the importance of isolating individual behaviors to understand the contribution of parts to the overall function. If we had been able to isolate a single function, maybe by having the processor produce the same math operation every single step, then the lesioning experiments could have produced more meaningful results. However, the same problem exists in neuroscience. It is extremely difficult or technically impossible to produce behaviors that only require a single aspect of the brain.

Beyond behavioral choices, we have equivalent problems in neuroscience that make the interpretation of lesioning data complicated [36]. In many ways the chip can be lesioned in a cleaner way than the brain: we can individually abolish every single transistor (this is only now becoming possible with neurons in simple systems [37, 38]). Even without this problem, finding that a lesion in a given area abolishes a function is hard to interpret in terms of the role of the area for general computation. And this ignores the tremendous plasticity in neural systems which can allow regions to take over for damaged areas. In addition to the statistical problems that arise from multiple hypothesis testing, it is obvious that the “causal relationship” we are learning is incredibly superficial: a given transistor is obviously not specialized for Donkey Kong or Space Invaders.

**Fig 5.**
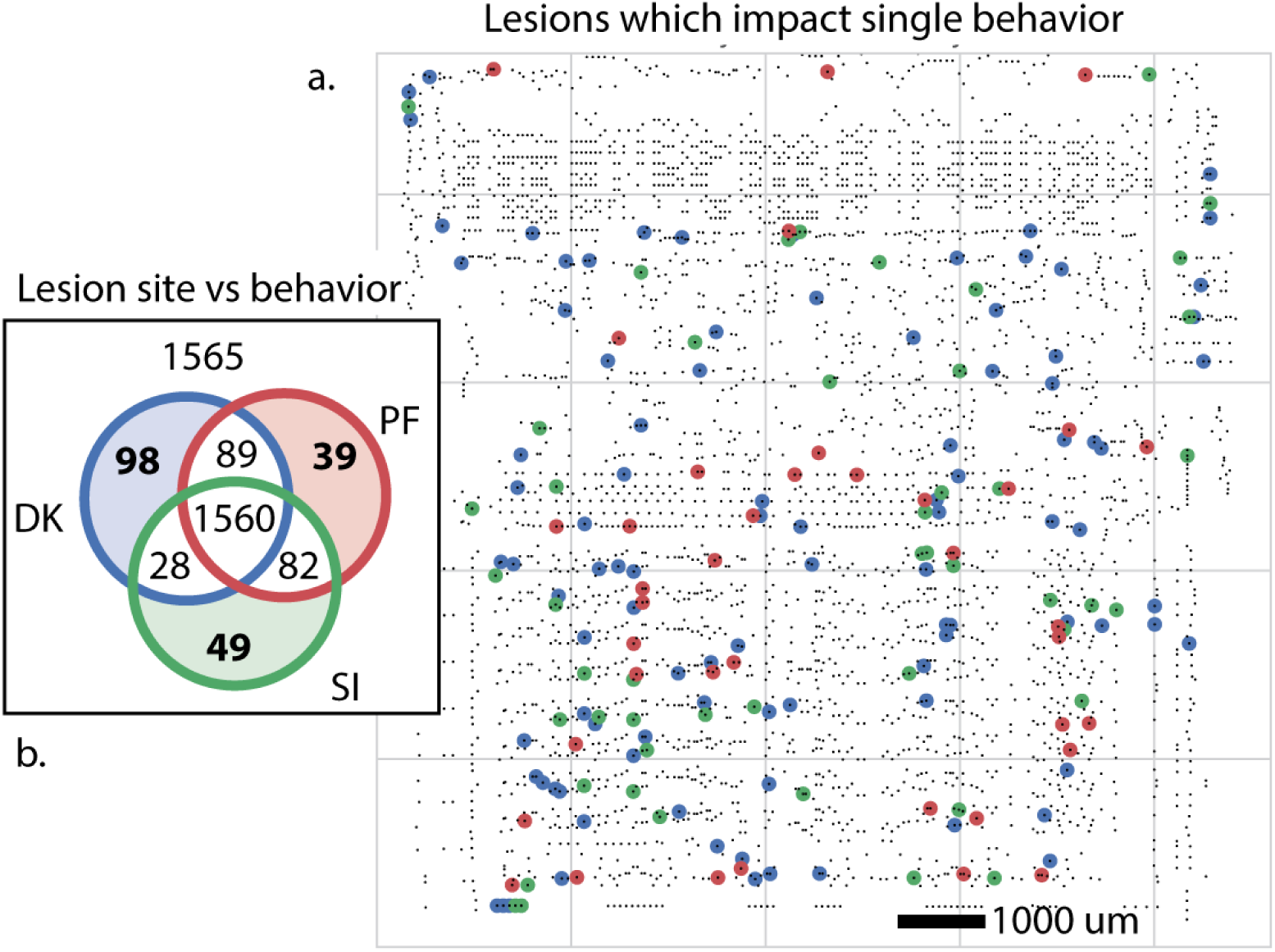
Lesioning every single transistor to identify function. We identify transistors whose elimination disrupts behavior analogous to lethal alleles or lesioned brain areas. These are transistors whose elimination results in the processor failing to render the game. (**A**) Transistors which impact only one behavior, colored by behavior. (**B**) Breakdown of the impact of transistor lesion by behavioral state. The elimination of 1565 transistors have no impact, and 1560 inhibit all behaviors.

**Fig 6.**
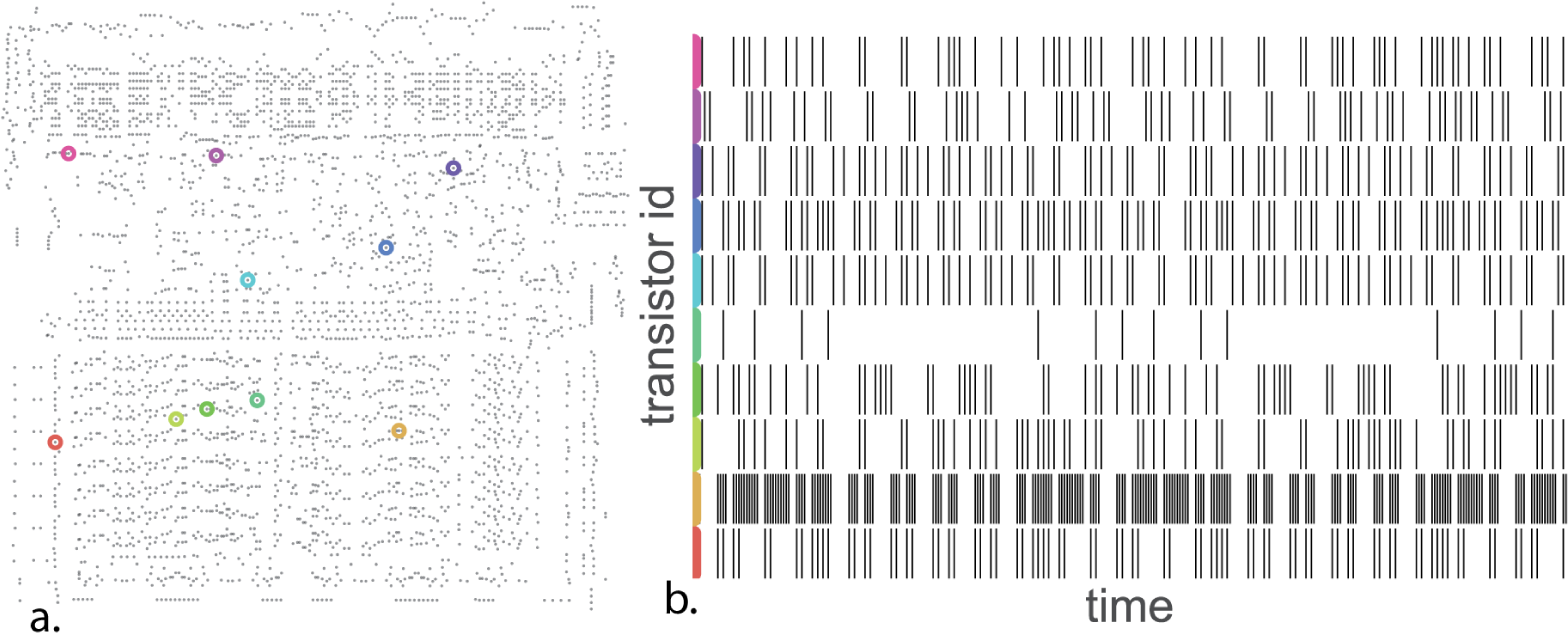
Analyzing the spikes to understand their statistics. (**A**) 10 identified transistors and (**B**) their spiking (rising edge) behavior over a short time window during behavior DK.

While in most organisms individual transistors are not vital, for many less-complex systems they are. Lesion individual interneurons in *C. elegans* or the H1 neuron in the fly can have marked behavioral impacts. And while lesioning larger pieces of circuitry, such as the entire TIA graphics chip, might allow for gross segregation of function, we take issue with this constituting “understanding”. Simply knowing functional localization, at any spatial scale, is only the most nacent step to the sorts of understanding we have outlined above.

## Analyzing tuning properties of individual transistors

We may want to try to understand the processor by understanding the activity of each individual transistor. We study the “off-to-on” transition, or “spike”, produced by each individual transistor. Each transistor will be activated at multiple points in time. Indeed, these transitions look surprisingly similar to the spike trains of neurons (fig 6). Following the standards in neuroscience we may then quantify the tuning selectivity of each transistor. For each of our transistors we can plot the spike rate as a function of the luminance of the most recently displayed pixel (fig 7). For a small number of transistors we find a strong tuning to the luminance of the most recently displayed pixel, which we can classify into simple (fig 7a) and (fig 7b) complex curves. Interestingly, however, we know for each of the five displayed transistors that they are not directly related to the luminance of the pixel to be written, despite their strong tuning. The transistors relate in a highly nonlinear way to the ultimate brightness of the screen. As such their apparent tuning is not really insightful about their role. In our case, it probably is related to differences across game stages. In the brain a neuron can calculate something, or be upstream or downstream of the calculation and still show apparent tuning making the inference of a neurons role from observational data very difficult [39]. This shows how obtaining an understanding of the processor from tuning curves is difficult.

**Fig 7.**
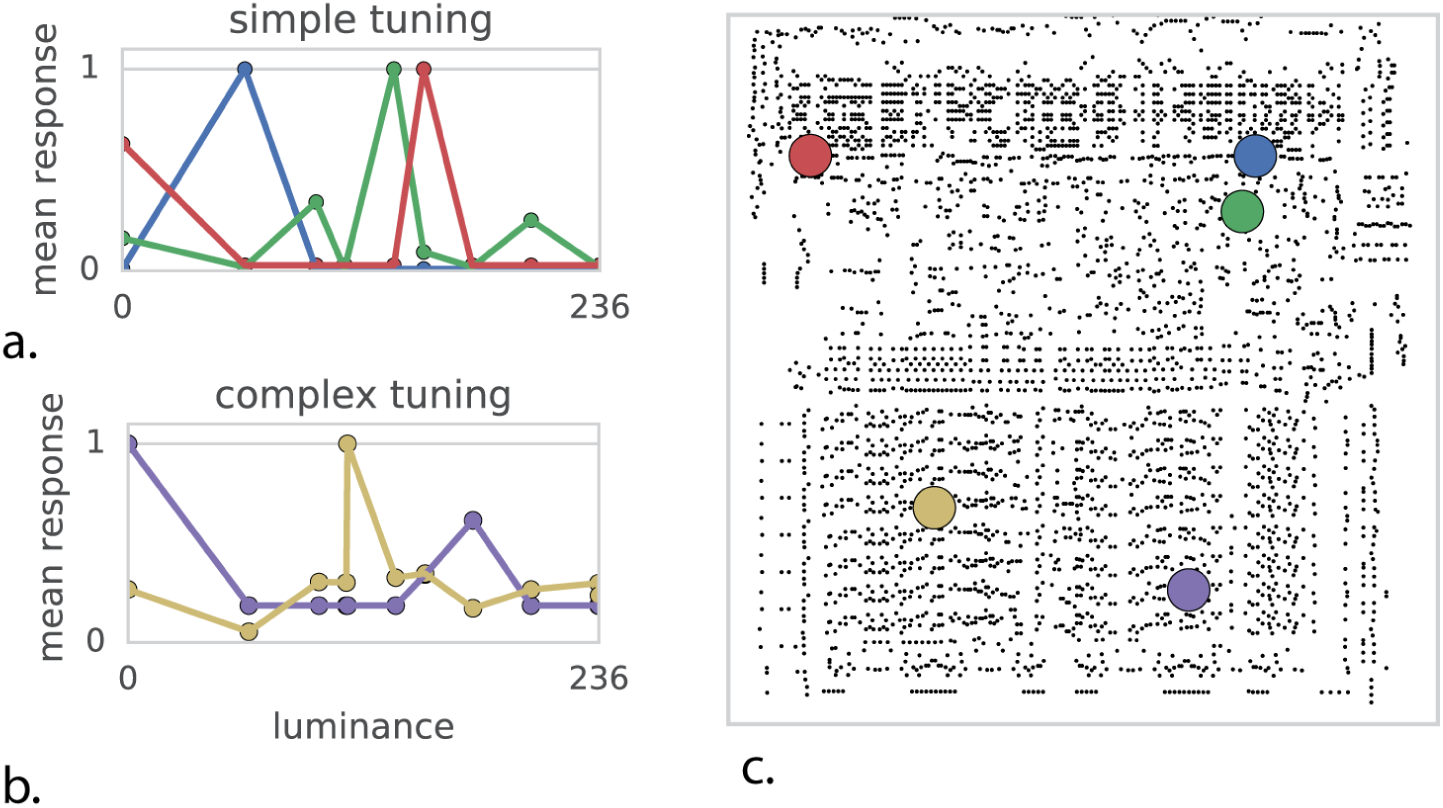
Quantifying tuning curves to understand function. Mean transistor response as a function of output pixel luminance. (**A**) Some transistors exhibit simple unimodal tuning curves. (**B**) More complex tuning curves. (**C**) Transistor location on chip.

Much of neuroscience is focused on understanding tuning properties of neurons, circuits, and brain areas [40–43]. Arguably this approach is more justified for the nervous system because brain areas are more strongly modular. However, this may well be an illusion and many studies that have looked carefully at brain areas have revealed a dazzling heterogeneity of responses [44–46]. Even if brain areas are grouped by function, examining the individual units within may not allow for conclusive insight into the nature of computation.

## The correlational structure exhibits weak pairwise and strong global correlations

Moving beyond correlating single units with behavior, we can examine the correlations present between individual transistors. We thus perform a spike-word analysis [47] by looking at “spike words” across 64 transistors in the processor. We find little to very weak correlation among most pairs of transistors (fig 8a). This weak correlation suggests modeling the transistors' activities as independent, but as we see from shuffle analysis (fig 8b), this assumption fails disastrously at predicting correlations across many transistors.

In neuroscience, it is known that pairwise correlations in neural systems can be incredibly weak, while still reflecting strong underlying coordinated activity. This is often assumed to lead to insights into the nature of interactions between neurons [47]. However, the processor has a very simple nature of interactions and yet produces remarkably similar spike word statistics. This again highlights how hard it is to derive functional insights from activity data using standard measures.

**Fig 8.**
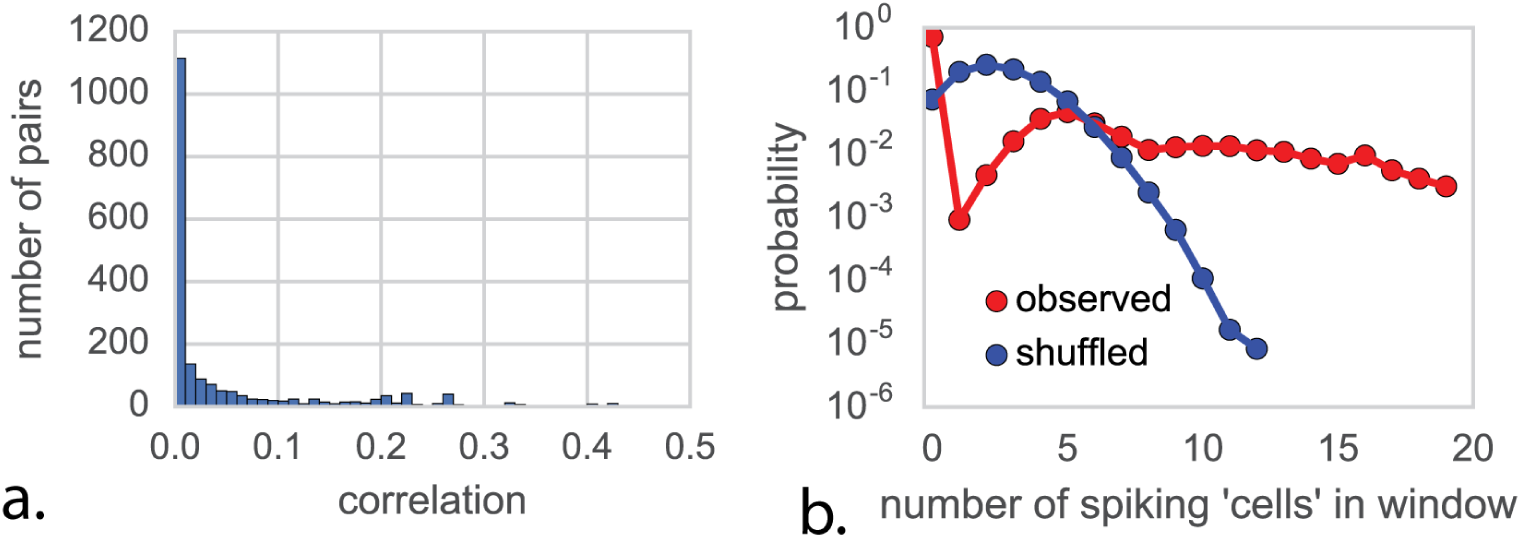
Spike-word analysis to understand synchronous states. (**A**) Pairs of transistors show very weak pairwise correlations during behavior SI, suggesting independence. (**B**) If transistors were independent, shuffling transistor labels (blue) would have no impact on the distribution of spikes per word, which is not the case (red)

**Fig 9.**
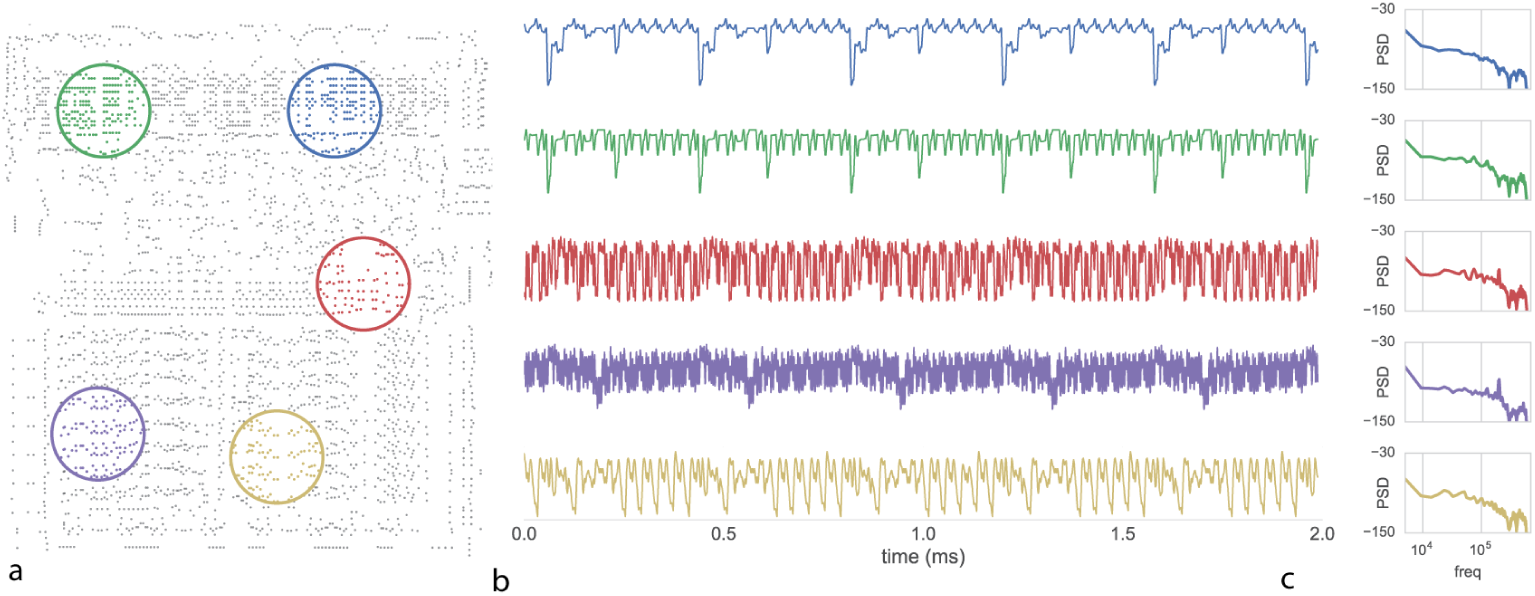
Examining local field potentials to understand network properties. We recorded from the processor during behavior DK. (**A**) Transistor switching is integrated and low-pass filtered over the indicated region. (**B**) local-field potential measurements from the indicated areas. (**C**) Spectral analysis of the indicated LFP regions identifies varying region-specific oscillations or “rhythms”

## Analyzing local field potentials

The activity of the entire chip may be high dimensional, yet we know that the chip, just like the brain, has some functional modularity. As such, we may be able to understand aspects of its function by analyzing the average activity within localized regions, in a way analogous to the local field potentials or the BOLD signals from functional magnetic imaging that are used in neuroscience. We thus analyzed data in spatially localized areas (fig 9a). Interestingly, these average activities look quite a bit like real brain signals (Fig 9b). Indeed, they show a rather similar frequency power relation of roughly power-law behavior. This is often seen as a strong sign of self-organized criticality [48]. Spectral analysis of the time-series reveals region-specific oscillations or “rhythms” that have been suggested to provide a clue to both local computation and overall inter-region communication. In the chip we know that while the oscillations may reflect underlying periodicity of activity, the specific frequencies and locations are epiphenomena. They arise as an artifact of the computation and tell us little about the underlying flow of information. And it is very hard to attribute (self-organized) criticality to the processor.

In neuroscience there is a rich tradition of analyzing the rhythms in brain regions, the distribution of power across frequencies as a function of the task, and the relation of oscillatory activity across space and time. However, the example of the processor shows that the relation of such measures to underlying function can be extremely complicated. In fact, the authors of this paper would have expected far more peaked frequency distributions for the chip. Moreover, the distribution of frequencies in the brain is often seen as indicative about the underlying biophysics. In our case, there is only one element, the transistor, and not multiple neurotransmitters. And yet, we see a similarly rich distribution of power in the frequency domain. This shows that complex multi-frequency behavior can emerge from the combination of many simple elements. Analyzing the frequency spectra of artifacts thus leads us to be careful about the interpretation of those occurring in the brain. Modeling the processor as a bunch of coupled oscillators, as is common in neuroscience, would make little sense.

## Granger causality to describe functional connectivity

Granger causality [49] has emerged as a method of assessing putative causal relationships between brain regions based on LFP data. Granger causality assesses the relationship between two timeseries *X* and *Y* by comparing the predictive power of two different time-series models to predict future values of *Y*. The first model uses only past values of *Y*, whereas the second uses the history of *X* and *Y*. The additon of *X* allows one to assess the putative “causality” (really, the predictive power) of *X*.

To see if we can understand information transmission pathways in the chip based on such techniques, we perform conditional Granger causality analysis on the above-indicated LFP regions for all three behavioral tasks, and plot the resulting inferences of causal interactions (fig 10). We find that the decoders affect the status bits. We also find that the registers are affected by the decoder, and that the accumulator is affected by the registers. We also find communication between the two parts of the decoder for Donkey Kong, and a lack of communication from the accumulator to the registers in Pitfall. Some of these findings are true, registers really affect the accumulator and decoders really affect the status bits. Other insights are less true, e.g. decoding is independent and the accumulator obviously affects the registers. While some high level insights may be possible, the insight into the actual function of the processor is limited.

The analysis that we did is very similar to the situation in neuroscience. In neuroscience as well, the signals come from a number of local sources. Moreover, there are also lots of connections but we hope that the methods will inform us about the relevant ones. It is hard to interpret the results - what exactly does the Granger causality model tell us about. Granger causality tells us how activity in the past are predictive of activity in the future, and the link from there to causal interactions is tentative at best [50] and yet such methods are extensively used across large subfields of neuroscience. Even if such methods would reliably tell us about large scale influences, it is very hard to get from a coarse resolution network to the microscopic computations.

**Fig 10.**
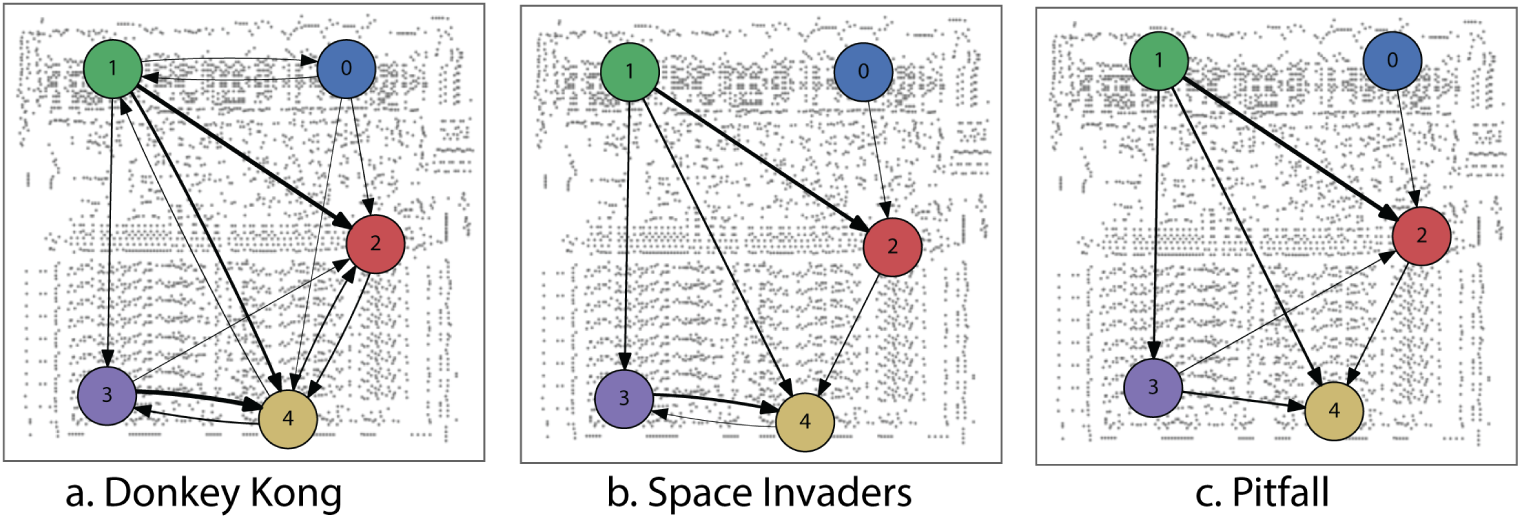
Analyzing conditional Granger causality to understand functional connectivity. Each of the recordings come from a well defined functional subcircuit. Green and blue are two parts of the decoder circuit. Red includes the status bits. Violet are part of the registers and yellow includes parts of the accumulator. We estimated for each behavioral state from LFP sites indicated in figure 9. Arrows indicate direction of Granger-causal relationship, arrow thickness indicates effect magnitude.

## Dimensionality reduction reveals global dynamics independent of behavior

In line with recent advances in whole-animal recordings [?, 2, 6, 7], we measure the activity across all 3510 transistors simultaneously for all three behavioral states (fig 11) and plot normalized activity for each transistor versus time. Much as in neural systems, some transistors are relatively quiet and some are quite active, with a clear behaviorally-specific periodicity visible in overall activity.

While whole-brain recording may facilitate identification of putative areas involved in particular behaviors [51], ultimately the spike-level activity at this scale is difficult to interpret. Thus scientists turn to dimensionality reduction techniques [2, 52, 53], which seek to explain high-dimensional data in terms of a low-dimensional representation of state. We use non-negative matrix factorization [54] to identify constituent signal parts across all time-varying transistor activity. We are thus, for the first time in the paper, taking advantage of all transistors simultaneously.

Non-negative matrix factorization assumes each recovered timeseries of transistor activity is a linear combination of a small number of underlying nonnegative time-varying signals (dimensions). Analogous with [2] we plot the recovered dimensions as a function of time (fig 12a) and the transistor activity profile of each component (fig 12b). We can also examine a map of transistor-component activity both statically (fig 12c) and dynamically (videos available in online supplementary materials). Clearly there is a lot of structure in this spatiotemporal dataset.

To derive insight into recovered dimensions, we can try and relate parts of the low-dimensional time series to known signals or variables we know are important (fig 13a). Indeed, we find that some components relate to both the onset and offset (rise and fall) of the clock signal(fig 13b,c). This is quite interesting as we know that the processor uses a two-phase clock. We also find that a component relates strongly to the processors read-write signal (fig 13d). Thus, we find that variables of interest are indeed encoded by the population activity in the processor.

**Fig 11.**
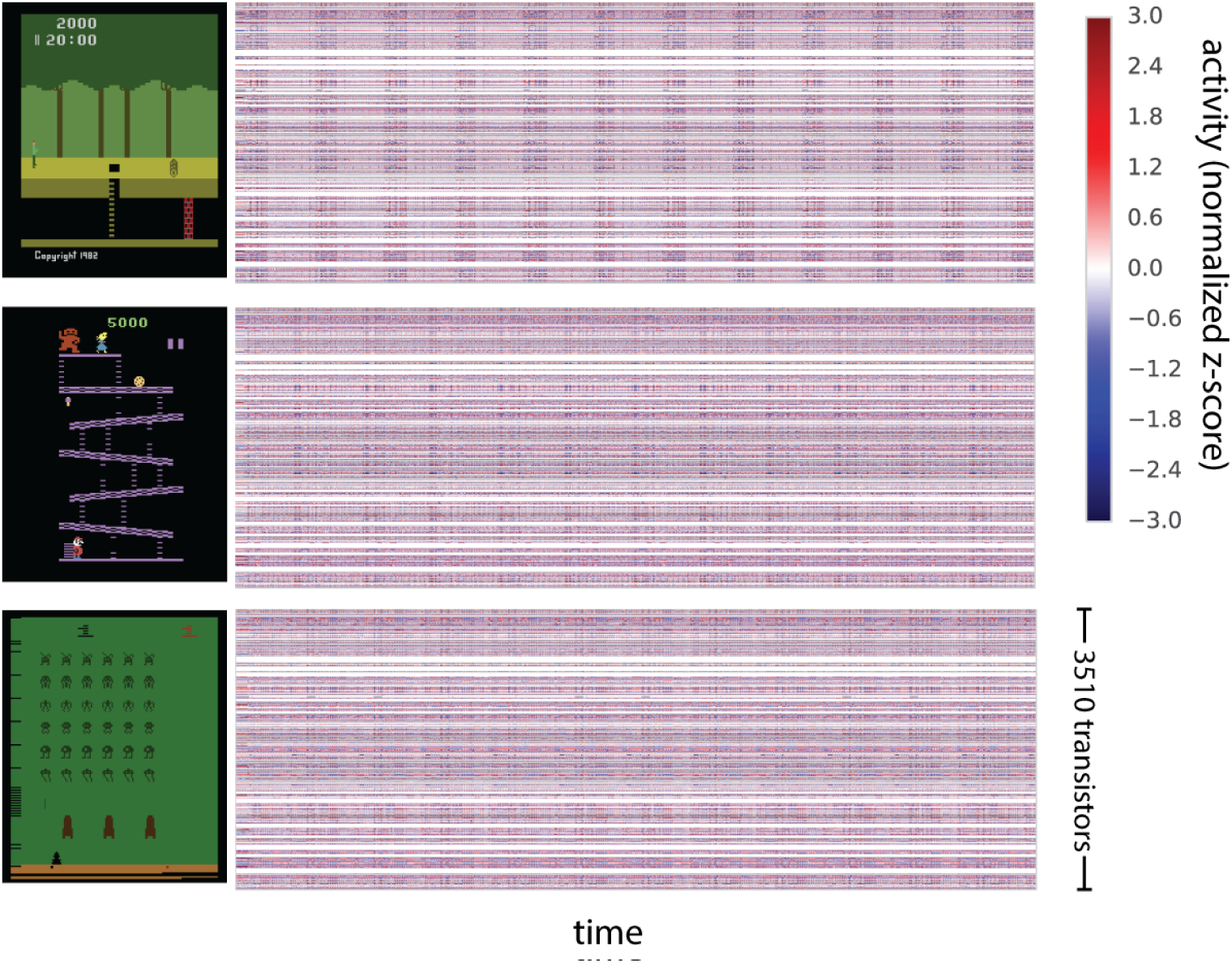
The processor activity map. For each of three behavioral states we plotted all the activities. Each transistor's activity is normalized to zero-mean and unit variance and plotted as a function of time.

**Fig 12.**
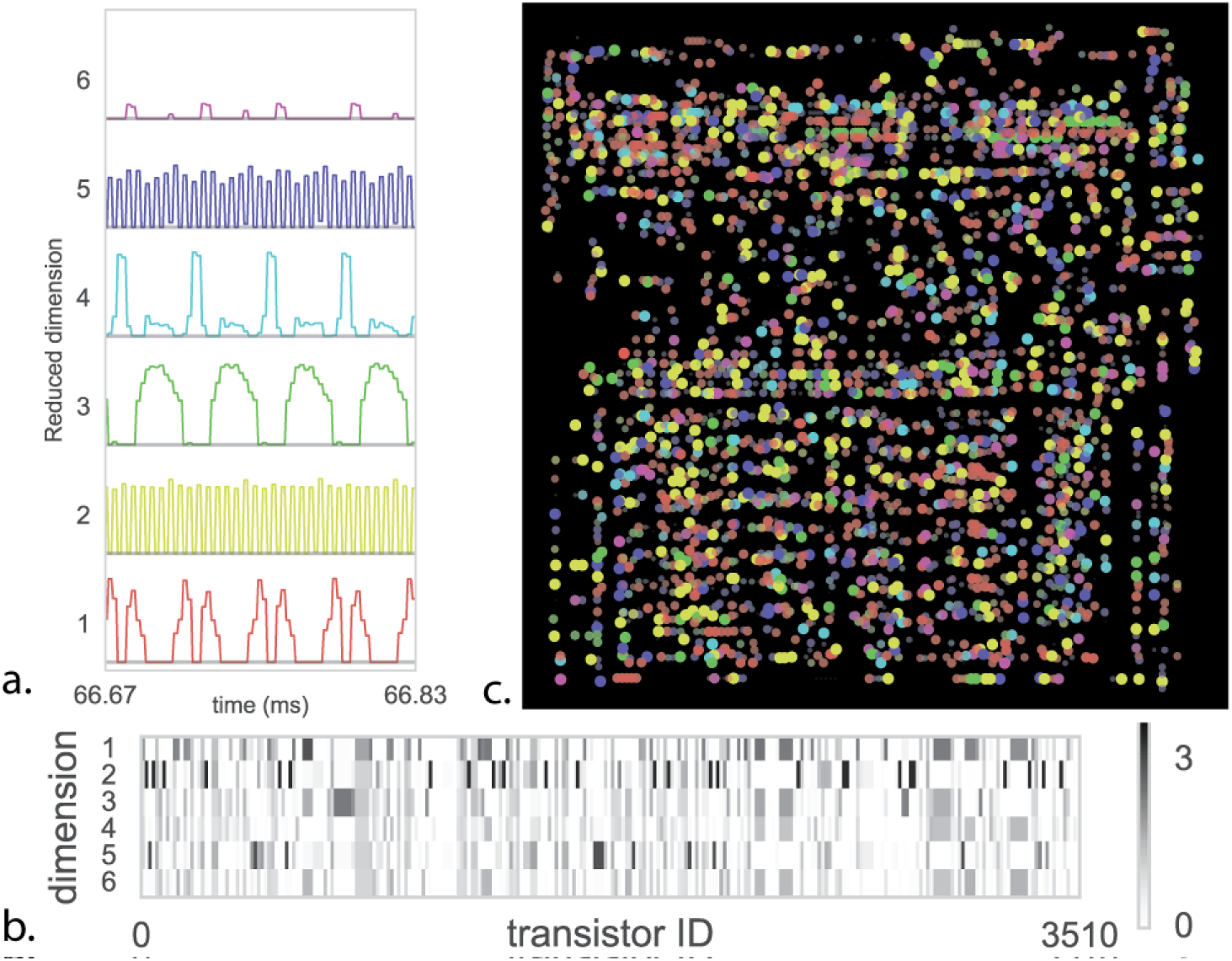
Dimensionality Reduction to understand the roles of transistors. We apply non-negative matrix factorization (NMF) to the space invaders (SI) task. (**A**) shows the six reduced dimensions as a function of time showing clear stereotyped activity. (**B**) the learned transistor state vectors for each dimension (**C**) Map of total activity — color indicates the dimension where the transistor has maximum value, and both saturation and point size indicate the magnitude of that value.

In neuroscience, it is also frequently found that components from dimensionality reduction relate to variables of interest [55, 56]. This is usually then seen as an indication that the brain cares about these variables. However, clearly, the link to the read-write signal and the clock does not lead to an overly important insight into the way the processor actually processes information. Similar questions arise in neuroscience where scientists ask if signals, such as synchrony, are a central part of information processing or if they are an irrelevant byproduct [57]. We should be careful at evaluating how much we understand and how much we are aided by more data.

Pondering the results of the processor analysis we can obtain some insights into the developments needed to better utilize dimensionality reduction towards an understanding. The narrow range of games that we considered and the narrow range of their internal states (we just simulated booting), means that many aspects of computation will not be reflected by the activities and hence not in the dimensionality reduction results. Moreover, the fact that we used linear reduction only allows for linear dependencies and transistors, just like neurons, have important nonlinear dependencies. Lastly, there is clearly a hierarchy in function in the processor and we would need to do it justice using hierarchical analysis approaches. The results of dimensionality reduction should be meaningful for guiding new experiments, necessitating transfer across chips in the same way as neuroscience experiments should transfer across animals. Importantly, the chip can work as a test case while we develop such methods.

## Discussion

Here we have taken a reconstructed and simulated processor and treated the data “recorded” from it in the same way we have been trained to analyze brain data. We have used it as a test case to check the na•ve use of various approaches used in neuroscience. We have found that the standard data analysis techniques produce results that are surprisingly similar to the results found about real brains. However, in the case of the processor we know its function and structure and our results stayed well short of what we would call a satisfying understanding.

Obviously the brain is not a processor, and a tremendous amount of effort and time have been spent characterizing these differences over the past century [21, 22, 58]. Neural systems are analog and and biophysically complex, they operate at temporal scales vastly slower than this classical processor but with far greater parallelism than is available in state of the art processors. Typical neurons also have several orders of magnitude more inputs than a transistor. Moreover, the design process for the brain (evolution) is dramatically different from that of the processor (the MOS6502 was designed by a small team of people over a few years). As such, we should be skeptical about generalizing from processors to the brain.

However, we cannot write off the failure of the methods we used on the processor simply because processors are different from neural systems. After all, the brain also consists of a large number of modules that can equally switch their input and output properties. It also has prominent oscillations, which may act as clock signals as well [59]. Similarly, a small number of relevant connections can produce drivers that are more important than those of the bulk of the activity. Also, the localization of function that is often assumed to simplify models of the brain is only a very rough approximation. This is true even in an area like V1 where a great diversity of co-localized cells can be found [60]. Altogether, there seems to be little reason to assume that any of the methods we used should be more meaningful on brains than on the processor.

**Fig 13.**
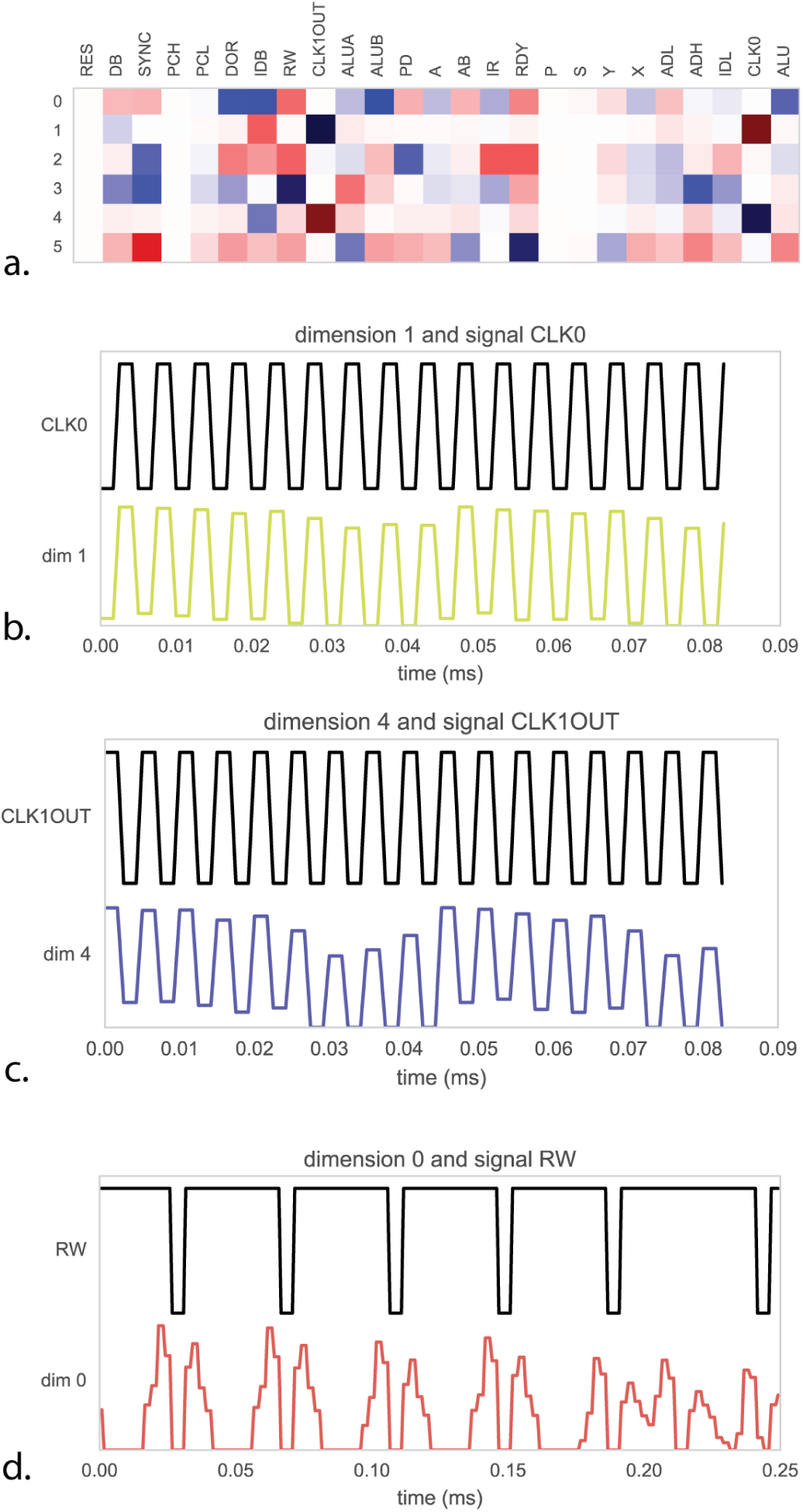
Relating dimensions to known signals to understanding the population code. (**A**) For each of the recovered dimensions in figure 12 we compute the correlation in time with 25 known signals inside the process. As we know the purpose of these signals we can measure how well the dimensions explain true underlying function. (**B**) Dimension 1 is strongly correlated with the processor clock CLK0, whereas (**C**) dimension 4 is correlated with the 180-degree out of phase CLK1OUT signal. (**D**) dimension 0 is strongly correlated with signal RW, indicating the processor switching between reading and writing memory.

To analyze our simulations we needed to convert the binary transistor state of the processor into spike trains so that we could apply methods from neuroscience to (see Methods). While this may be artefactual, we want to remind the reader that in neuroscience the idea of an action potential is also only an approximate description of the effects of a cell's activity. For example, there are known effects based on the extrasynaptic diffusion of neurotransmitters [61] and it is believed that active conductances in dendrites may be crucial to computation [62].

Our behavioral mechanisms are entirely passive as both the transistor based simulator is too slow to play the game for any reasonable duration and the hardware for game input/output has yet to be reconstructed. Even if we could “play” the game, the dimensionality of the input space would consist at best of a few digital switches and a simple joystick. One is reminded of the reaching tasks which dominate a large fraction of movement research. Tasks that isolate one kind of computation would be needed so that interference studies would be really interpretable.

If we had a way of hypothesizing the right structure, then it would be reasonably easy to test. Indeed, there are a number of large scale theories of the brain [5, 63, 64]. However, the set of potential models of the brain is unbelievably large. Our data about the brain from all the experiments so far, is very limited and based on the techniques that we reviewed above. As such, it would be quite impressive if any of these high level models would actually match the human brain to a reasonable degree. Still, they provide beautiful inspiration for a lot of ongoing neuroscience research and are starting to exhibit some human-like behaviors [63]. If the brain is actually simple, then a human can guess a model, and through hypothesis generation and falsification we may eventually obtain that model. If the brain is not actually simple, then this approach may not ever converge. Simpler models might yield more insight – specifically seeking out an “adder” circuit might be possible, if we had a strong understanding of binary encoding and could tease apart the system to specifically control inputs and outputs of a subregion – examine it in slice, if you will.

The analytic tools we have adopted are in many ways “classic”, and are taught to graduate students in neuroinformatics courses. Recent progress in methods for dimensionality reduction, subspace identification, time-series analysis, and tools for building rich probabilistic models may provide some additional insight, assuming the challenges of scale can be overcome. Culturally, applying these methods to real data, and rewarding those who innovate methodologically, may become more important. We can look at the rise of bioinformatics as an independent field with its own funding streams. Neuroscience needs strong neuroinformatics to make sense of the emerging datasets and known artificial systems can serve as a sanity check and a way of understanding failure modes.

We also want to suggest that it may be an important intermediate step for neuroscience to develop methods that allow understanding a processor. Because they can be simulated in any computer and arbitrarily perturbed, they are a great testbed to ask how useful the methods are that we are using in neuroscience on a daily basis. Scientific fields often work well in situations where we can measure how well a project is doing. In the case of processors we know their function and we can know if our algorithms discover it. Unless our methods can deal with a simple processor, how could we expect it to work on our own brain? Machine learning and statistics currently lack good high-dimensional datasets with complex underlying dynamics and known ground truth. While not a perfect match, the dynamics of a processor may provide a compelling intermediate step. Additionally, most neural datasets are still “small data” – hundreds of cells over tens of minutes. The processor enables the generation of arbitrary complexity and arbitrarially-long timeseries, enabling a focus on *scalable* algorithms. We must be careful to not over-fit, but neuroscience is rife with examples of adopting analytic tools from vary different domains (linear system theory, stochastic process theory, kalman filtering) to understand neural systems.

**Fig 14.**
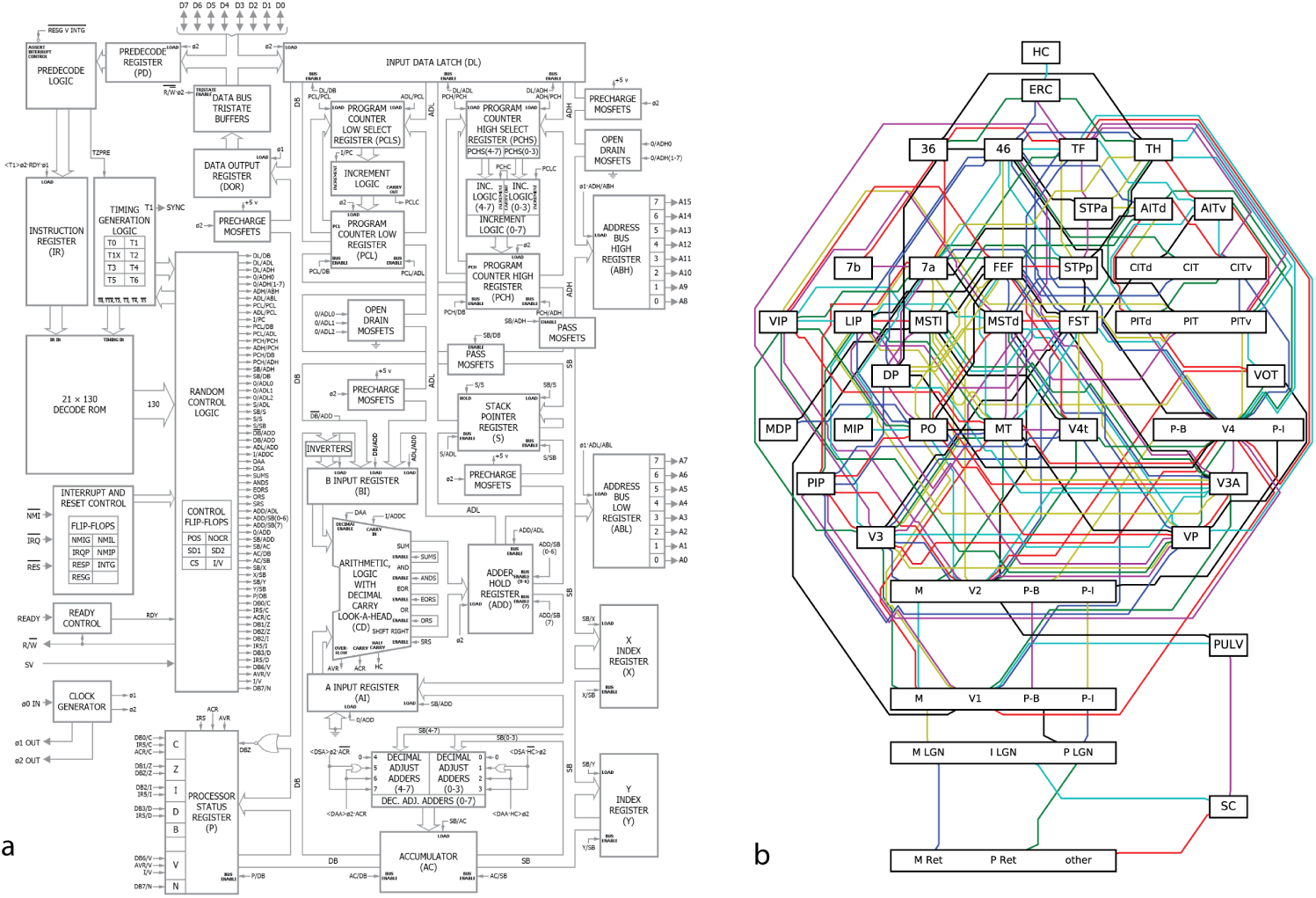
Understanding the processor. (**A**) For the processor we know which part of the the chip is responsible for which function. We know that these are meaningful because the designers told us so. And for each of these modules we know how the outputs depend on the inputs. (**B**) For the brain, it is harder to be sure. The primate visual system is often depicted in a similar way, such as this diagram adapted from the classic Felleman and vanEssen [65] diagram. These areas are primarially divided according to anatomy, but there is extensive debate about the ideal way of dividing the brain into functional areas. Moreover, we currently have little of an understanding how each area's outputs depend on its inputs.

In the case of the processor, we really understand how it works. We have a name for each of the modules on the chip and we know which area is covered by each of them (fig 14a). Moreover, for each of these modules we know how its outputs depend on its inputs and many students of electrical engineering would know multiple ways of implementing the same function. In the case of the brain, we also have a way of dividing it into regions (fig 14b). However, we only use anatomy to divide into modules and even among specialists there is a lot of disagreement about the division. Most importantly though, we do not generally know how the output relates to the inputs. As we reviewed in this paper, we may even want to be careful about the conclusions about the modules that neuroscience has drawn so far, after all, much of our insights come from small datasets, with analysis methods that make questionable assumptions.

There are other computing systems that scientists are trying to reverse engineer. One particularly relevant one are artificial neural networks. A plethora of methods are being developed to ask how they work. This includes ways of letting the networks paint images [66] and ways of plotting the optimal stimuli for various areas [67]. While progress has been made on understanding the mechanisms and architecture for networks performing image classification, more complex systems are still completely opaque [68]. Thus a true understanding even for these comparatively simple, human-engineered systems remains elusive, and sometimes they can even surprise us by having truly surprising properties [69]. The brain is clearly far more complicated and our difficulty at understanding deep learning may suggest that the brain is hard to understand if it uses anything like gradient descent on a cost function.

What kind of developments would make understanding the processor, and ultimately the brain, more tractable? While we can offer no definitive conclusion, we see multiple ways in which we could have better understood the processor. If we had experiments that would more cleanly separate one computation then results would be more meaningful. For example, lesion studies would be far more meaningful if we could also simultaneously control the exact code the processor was executing at a given moment. Better theories could most obviously have helped; if we had known that the microprocessor has adders we could have searched for them. Lastly, better data analysis methods, e.g. those that can explicitly search for hierarchical structure or utilize information across multiple processors. Development in these areas seems particularly promising. The microprocessor may help us by being a sieve for ideas: good ideas for understanding the brain should also help us understand the processor. Ultimately, the problem is not that neuroscientists could not understand a microprocessor, the problem is that they would not understand it given the approaches they are currently taking.

## Methods

### Netlist acquisition

All acquisition and development of the initial simulation was performed in James [10]. 200° F sulfuric acid was used to decap multiple 6502D ICs. Nikon LV150n and Nikon Optiphot 220 light microscopes were used to capture 72 tiled visible-light images of the die, resulting in 342 Mpix of data. Computational methods and human manual annotation used developed to reconstruct the metal, polysilicon, via, and interconnect layers. 3510 active enhancement-mode transistors were captured this way. The authors inferred 1018 depletion-mode transistors (serving as pullups) from the circuit topology as they were unable to capture the depletion mask layer.

### Simulation and behaviors

An optimized C++ simulator was constructed to enable simulation at the rate of 1000 processor clock cycles per wallclock second. We evaluated the four provided ROMs (Donkey Kong, Space Invaders, Pitfall, and Asteroids) ultimately choosing the first three as they reliably drove the TIA and subsequently produced image frames. 10 seconds of behavior were simulated for each game, resulting in over 250 frames per game.

### Lesion studies

Whole-circuit simulation enables high-throughput targeted manipulation of the underlying circuit. We systematically perturb each transistor in the processor by forcing its input high, thus leaving it in an “on” state. We measure the impact of a lesion by whether or not the system advances far enough to draw the first frame of the game. Failure to produce the first frame constitutes as a loss of function. We identified 1560 transistors which resulted in loss of function across all games, 200 transistors which resulted in loss of function across two games, and 186 transistors which resulted in loss of function for a single game. We plot those single-behavior lesion transistors by game in figure 5.

### Connectomic Analysis

Using the acquired netlist, we implement the authors method from [29] on the region of the processor consisting of the X, Y, and S registers. A nonparametric distance-dependent stochastic block model is jointly fit to six connectivitiy matrices : *G* → *C*1, *G* → *C*2, *C*1 → *C*2 *C*2 → *C*1, *C*1 → *G, C*2 → *G*, and via Markov-chain Monte Carlo, seeks the maximum a posteriori estmate for the observed connectivity.

### Spiking

We chose to focus on transistor switching as this is the closest in spirit to discrete action potentials of the sort readily available to neuroscientific analysis. The alternative, performing analysis with the signals on internal wires, would be analogous to measuring transmembrane voltage. Rasters were plotted from 10 example transistors which showed sufficient variance in spiking rate.

### Tuning curves

We compute luminance from the RGB output value of the simulator for each output pixel to the TIA. We then look at the transistor rasters and sum activity for 100 previous timesteps and call this the “mean rate”. For each transistor we then compute a tuning curve of mean rate versus luminance, normalized by the frequency of occurrence of that luminance value. Note that each game outputs only a small number of discrete colors and thus discrete luminance values. We used SI as it gave the most equal sampling of luminance space. We then evaluate the degree of fit to a unimodial Gaussian for each resulting tuning curve and classify tuning curves by eye into simple and complex responses, of which figure 5 contains representative examples.

### Spike-word analysis

For the SI behavior we took spiking activity from the first 100ms of SI and performed spike word analysis on a random subset of 64 transistors close to the mean firing rate of all 3510.

### Local Field Potential

To derive “local field potentials” we spatially integrate transistor switching over a region with a Gaussian weighting of *σ* = 500*μm* and low-pass filter the result using a window with a width of 4 timesteps.

We compute periodograms using Welch's method with 256-sample long windows with no overlap and a Hanning window.

### Granger Causality

We adopt methods for assessing conditional Granger causality as outlined in [70]. We take the LFP generated using methods in section and create 100 1ms-long trials for each behavioral experiment. We then compute the conditional Granger causality for model orders ranging from 1 to 31. We compute BIC for all behaviors and select a model order of 20 as this is where BIC plateaus.

### Whole brain recording

The transistor switching state for the first 10^6^ timestamps for each behavioral state is acquired, and binned in 100-timestep increments. The activity of each transistor is converted into a z-score by subtracting mean and normalizing to unit variance.

### Dimensionality Reduction

We perform dimensionality reduction on the first 100,000 timesteps of the 3510-element transistor state vectors for each behavioral condition. We use non-negative matrix factorization, which attempts to find two matrices, *W* and *H*, whose product *W H* approximates the observed data matrix *X*. This is equivalent to minimizing the objective 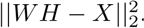.

The Scikit-Learn [71] implementation initialized via nonnegative double singular value decomposition solved via coordinate descent, as is the default. We use a latent dimensionality of 6 as it was found by hand to provide the most interpretable results. When plotting, the intensity of each transistor in a latent dimension is indicated by the saturation and size of point.

To interpret the latent structure we first compute the signed correlation between the latent dimension and each of the 25 known signals. We show particularly interpretable results.

## Acknowledgments

We'd like to thank the Visual 6502 team for the original simulation and reconstruction work. We thank Gary Marcus, Adam Marblestone, Malcolm MacIver, John Krakauer, and Yarden Katz for helpful discussions, and The Kavli Foundation for sponsoring the “Workshop on Cortical Computation” where these ideas were first developed. Thanks to Phil Mainwaring for providing the schematic of the 6502 in fig 14. EJ is supported in part by NSF CISE Expeditions Award CCF-1139158, DOE Award SN10040 DE-SC0012463, and DARPA XData Award FA8750-12-2-0331, and gifts from Amazon Web Services, Google, IBM, SAP, The Thomas and Stacey Siebel Foundation, Adatao, Adobe, Apple, Inc., Blue Goji, Bosch, Cisco, Cray, Cloudera, EMC2, Ericsson, Facebook, Fujitsu, Guavus, HP, Huawei, Informatica, Intel, Microsoft, NetApp, Pivotal, Samsung, Schlumberger, Splunk, Virdata, and VMware. KPK is supported by the National Institutes of Health (MH103910, NS074044, EY021579).

